# Conservation genomics of an Australian orchid complex with implications for the taxonomic and conservation status of *Corybas dowlingii*

**DOI:** 10.1101/2020.01.22.916080

**Authors:** Natascha D. Wagner, Mark A. Clements, Lalita Simpson, Katharina Nargar

## Abstract

This study assessed genomic diversity in an Australian species complex in the helmet orchids to clarify taxonomic delimitation and conservation status of the threatened species *Corybas dowlingii,* a narrow endemic from southeast Australia. Taxonomic delimitation between the three closely related species *C. aconitiflorus*, *C. barbarae,* and *C. dowlingii* has been mainly based on floral traits which exhibit varying degrees of overlap, rendering species delimitation in the complex difficult. Genomic data for the species complex was generated using double-digest restriction-site associated DNA (ddRAD) sequencing. Maximum likelihood, NeighborNet, and Bayesian structure analyses showed genetic differentiation within the species complex and retrieved genomic signatures consistent with hybridisation and introgression between *C. aconitiflorus* and *C. barbarae,* and an intermediate genetic position of *C. dowlingii* indicating a hybrid origin of the species. The genetic structure analysis showed varying levels of genetic admixture for several *C. aconitiflorus*, *C. barbarae,* and *C. dowlingii* samples, thus further corroborating the presence of hybridisation and introgression within the species complex. The taxonomic status of *C. dowlingii* D.L.Jones was revised to *C. × dowlingii* D.L.Jones *stat. nov.* to reflect its hybrid origin. The conservation status of *C. × dowlingii* was assessed based on key ecological and ethical aspects, and recommendations made regarding its conservation status in Australian conservation legislation.

## Introduction

Orchidaceae is the world’s second largest flowering plant family, comprising over 27,800 species and 736 genera (Chase et al. 2015; (List 2018) that display extraordinary morphological and ecological diversity. Orchids inhabit a wide range of terrestrial and epiphytic habitats worldwide exhibiting their highest species diversity in the tropics (Cribb et al. 2003). However, many orchid species possess only narrow distributions, rendering them vulnerable to threats such as climate change, habitat destruction or over-exploitation (Cribb et al. 2003).

In the Australian flora, orchids rank within the ten largest angiosperm families, and exhibit a high level of endemicity with about 90 % of Australian native orchids found nowhere else (Govaerts et al. 2017). Orchids constitute over 15 % of Australia’s threatened plant species and over one third of Australia’s critically endangered plants (EPBC 1999). Over the last decade, new orchid species have been described at a rate of approximately 500 species per year worldwide (Chase et al. 2015). However, species delimitation in orchids is often challenging, which is reflected by the large number of taxonomic synonyms in orchids (Govaerts et al. 2017). In Australia, the number of orchid species recognised for the country has increased from around 900 species to more than 1,300 over the past two decades (Hopper 2009; Govaerts et al. 2017). This increase has been partly attributed to the discovery and description of new species and partly to the application of narrower species concepts elevating many infraspecific taxa to species level (Hopper 2009; Chase et al. 2015). The resulting uncertainties in the taxonomic status of many Australian orchid species greatly hamper effective conservation management and allocation of scarce resources.

The *Corybas aconitiflorus* complex (Acianthinae, Diurideae) is a group of small geophytic, colony-forming herbs with globose tubers. They possess a single heart-shaped leaf and a solitary flower with an enlarged hood-shaped dorsal sepal that imparts the impression of a helmet, hence the vernacular name ‘helmet orchids’ (Pridgeon et al. 2001). The flowers are mostly dull-coloured with shades of reddish or brownish purple, sometimes white to pinkish, and are pollinated by fungus gnats of the family Mycetophilidae Newman 1834 (Pridgeon et al. 2001). As in most orchids, the fruit is a dehiscent capsule with numerous dust-like seeds.

Previous molecular studies provided insights into phylogenetic relationships in *Corybas* (Clements et al. 2002, Lyon 2014) and identified an unresolved clade of four closely related species which constitute the *C. aconitiflorus* complex: three of the species are endemic to Australia (*C. aconitiflorus* Salisb., *C. barbarae* D.L.Jones, and *C. dowlingii* D.L.Jones) and one to Java (*C. imperiatorus* (J.J.Sm.) Schltr.) (Lyon 2014). Among the Australian species of the complex, *C. aconitiflorus* and *C. barbarae* are locally common and widespread, extending over 2,000 km along the Australian east coast, broadly in sympatry (AVH 2018). In contrast, *C. dowlingii* is narrowly endemic in New South Wales, extending ca. 100 km from Bulahdelah north of Newcastle to Freemans Waterhole south of Newcastle (Jones 2004; AVH 2018)), and occurs within the distribution range of both *C. aconitiflorus* and *C. barbarae* (AVH 2018). *Corybas dowlingii* is listed as vulnerable species at federal level (EPBC 1999) and as endangered under the New South Wales Threatened Species Conservation Act (1995) due to its highly restricted distribution and anthropogenic pressures on its habitat.

Previous molecular systematic studies in Diurideae based on the internal transcribed spacer (ITS) confirmed the taxonomic placement of *Corybas* within subtribe Acianthinae (Kores et al. 2001; Clements et al. 2002), and a combined phylogenetic analysis of ITS and 68 morphological characters in Acianthinae showed the *C. aconitiflorus* complex as unresolved polytomy together with *C. cerasinus,* an endemic species from northern Queensland (Australia) and New Zealand endemic *C. cheesemanii* (Clements et al. 2002). However, *C. dowlingii* was not described at the time of the study of (Clements et al. 2002). A molecular study in *Corybas* based on five plastid and three nuclear markers further resolved phylogenetic relationships in the genus and showed *C. aconitiflorus*, *C. barbarae*, *C. dowlingii*, and *C. imperatorius* as a clade, however relationships among these species remained unclear due to lack of statistical support (Lyon 2014). While sequence divergence among the three Australian species was shallow, *C. imperiosus* was situated on a long branch in the phylogenetic tree reconstruction (Lyon 2014). Morphologically, *C. dowlingii* is only weakly differentiated from *C. aconitiflorus* and *C. barbarae*. Its flowers are dark purplish red, and the labellum is of the same colour with some whitish areas and sparse bristles (Jones 2004, 2006). It is distinguished from the other two species mainly by differences in flower size and colouration, and a later, yet overlapping flowering time. *Corybas aconitiflorus* has greyish to reddish purple flowers that are smaller than in *C. dowlingii*, and has a whitish labellum with sparse tiny bristles (Jones 2004, 2006). *Corybas barbarae* has white to pinkish flowers of a similar size to *C. dowlingii* and possesses a white labellum covered in dense bristles (Jones 2004, 2006). Due to the low genetic divergence within the *C. aconitiflorus* complex observed in previous molecular studies (Clements et al. 2002; Lyon 2014), and the weak morphological differentiation between the three species with partly overlapping character states, further molecular studies with more informative molecular markers are required to assess species delimitation within the complex and to re-evaluate the conservation status of *C. dowlingii*.

Recent advances in high-throughput DNA sequencing and bioinformatics offer powerful genomic approaches to resolve complex inter- and intraspecific relationships at unprecedented resolution, facilitating the re-assessment of taxonomic concepts in species complexes and the conservation status of rare and threatened species (Ahrens et al. 2017; Bateman et al. 2018; Coates et al. 2018; Cozzolino et al. 2019; Taylor and Larson 2019).

Restriction-site-associated DNA sequencing (Baird et al. 2008) is a next generation sequencing method based on reduced representation library sequencing, which is a cost-effective method to obtain genome-scale data from non-model organisms (Davey et al. 2011; Lemmon and Lemmon 2013). For RADseq, genomic DNA is digested using one or two restriction enzymes (ddRAD). DNA fragments of a certain size range are selected as subset for library preparation and then subjected to high throughput sequencing (Miller et al. 2007; Baird et al. 2008; Peterson et al. 2012)(Miller et al. 2007; Lemmon and Lemmon 2013). RADseq has been successfully used to clarify inter- and intraspecific relationships and to assess taxonomic delimitation in species complexes, including in orchids (Brandrud et al., 2019a, b; Wagner et al. 2013; Jones et al. 2013; Eaton and Ree 2013; Escudero et al. 2014; Takahashi et al. 2014; Mort et al. 2015; Herrera and Shank 2016; Beheregaray et al. 2017; Bateman et al. 2018; Hipp et al. 2018).

The aims of this study were to assess genetic diversity and structure within the *C. aconitiflorus* complex based on genomic data derived from double-digest restriction-site associated DNA sequencing (ddRADseq) in order to clarify species delimitation within the complex and to evaluate the taxonomic and conservation status of the narrow endemic *C. dowlingii*.

## Materials and methods

### Material studied

In total, 72 samples were included in the study, of which 70 samples from the *C. aconitiflorus* complex: *C. aconitiflorus* (24 samples, 9 localities), *C. barbarae* (32 samples, 5 localities), and *C. dowlingii* (14 samples, 2 localities). *Corybas pruinosus* (A.Cunn.) Rchb.f. (2 samples) was included as outgroup based on Clements et al. (2002). Sampling focussed on the south-eastern distribution of the *C. aconitiflorus* complex to which *C. dowlingii* is endemic and extended from the restricted distribution of *C. dowlingii* (between Port Macquarie and Newcastle, New South Wales) ca. 300 km northwards to the border between New South Wales and Queensland (Uralba), ca. 1,200 km southwards to Tasmania (Ulverstone), and ca. 600 km eastwards to Lord Howe Island. For each population, one herbarium voucher was taken per species sampled and lodged at CANB. Sampling locations are shown in Fig. 1 and further details on plant material studied are provided in Tab. 1.

**Fig. 1.**
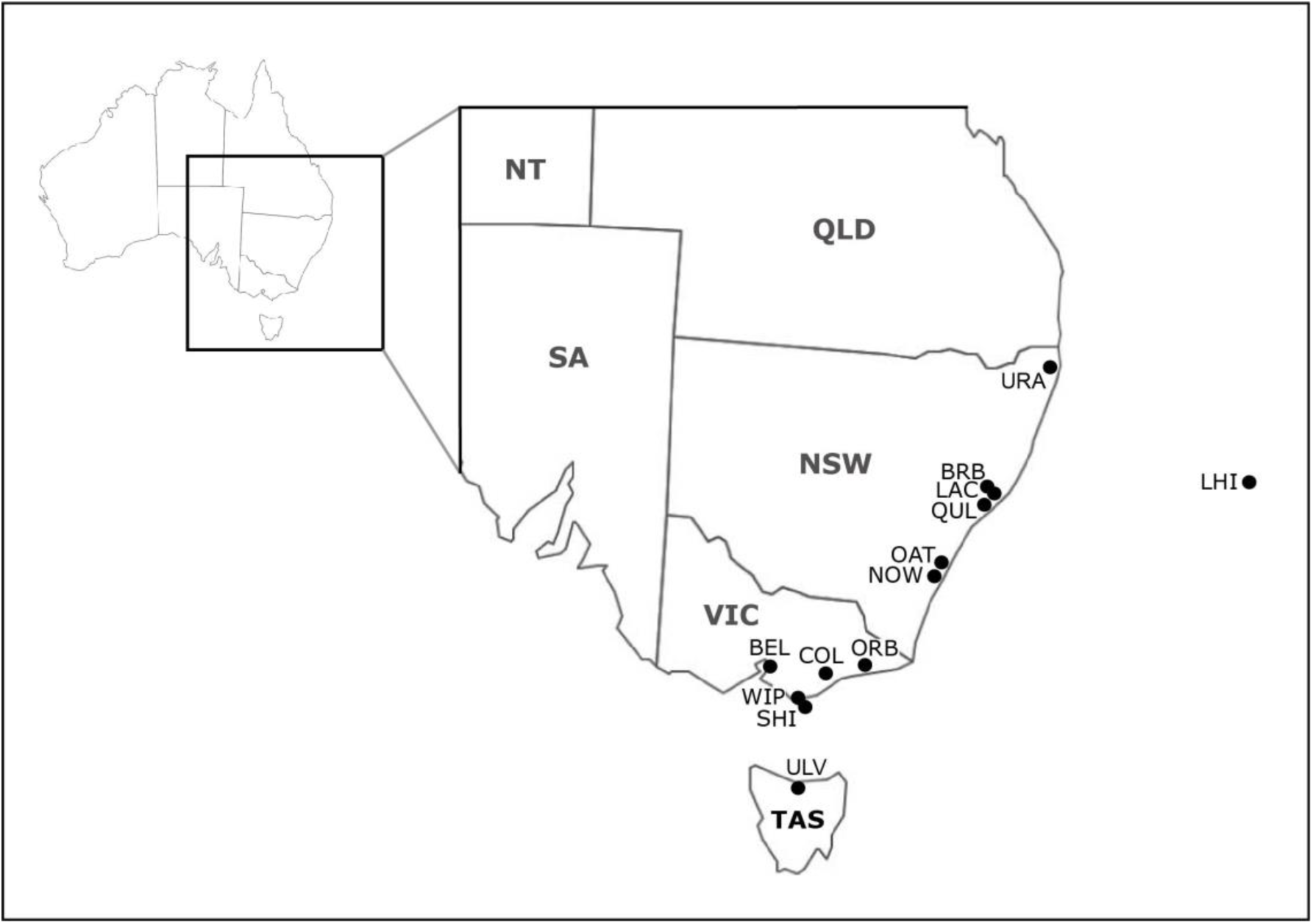
Sampling locations for study of genomic diversity of the *C. aconitiflorus* complex in southeastern Australia. Australian state abbreviations in bold. BEL: Belgrave South; BRB: Broken Bago; COL: Colquhoun; LAC: Lake Cathie; LHI: Lord Howe Island; NOW: Nowra; OAT: Oatly Park; ORB: Orbost; QUL: Queens Lake; SHI: Shallow Inlet; URA: Uralba Nature Reserve; ULV: Ulverstone; WIP: Wilson Promontory

**Tab. 1.**
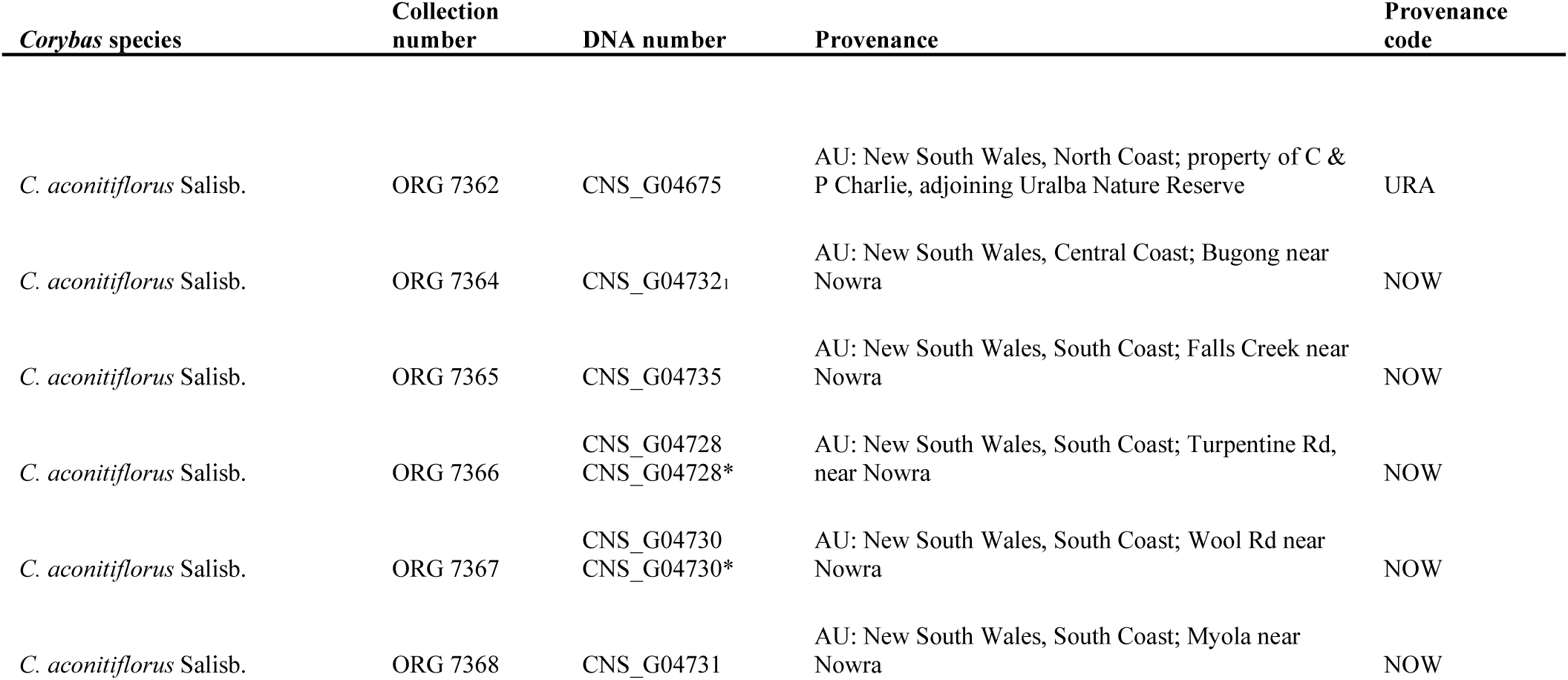
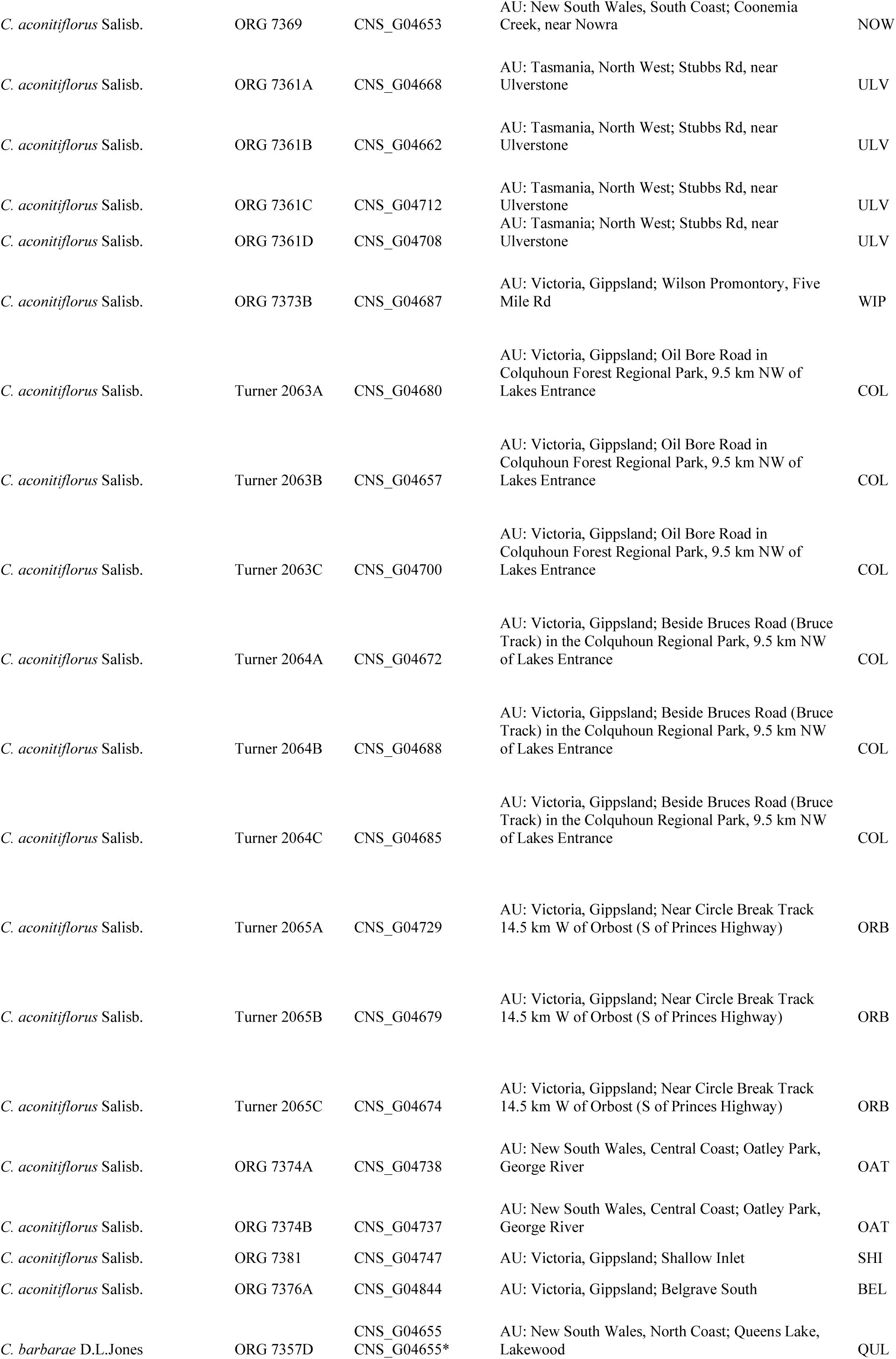
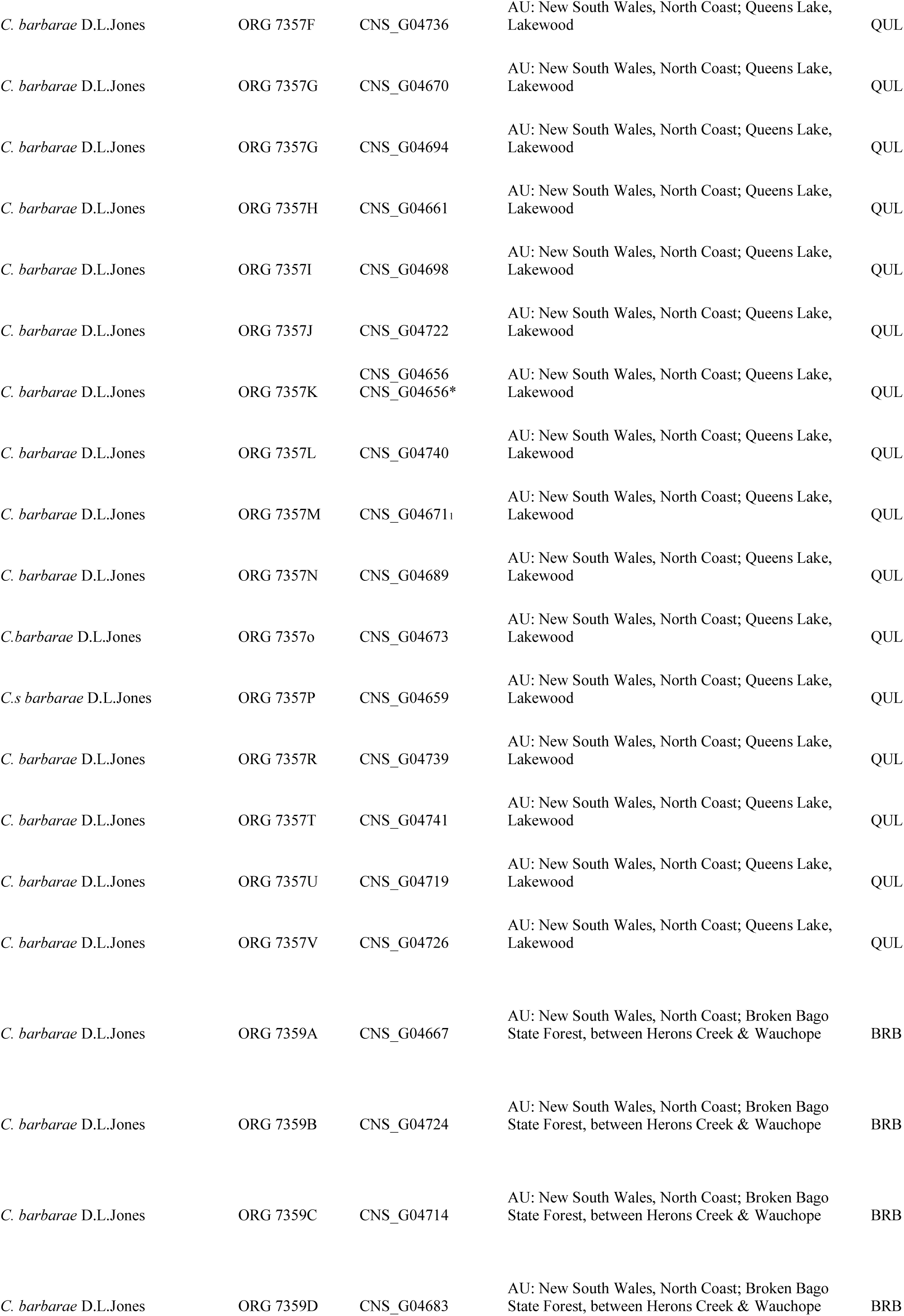
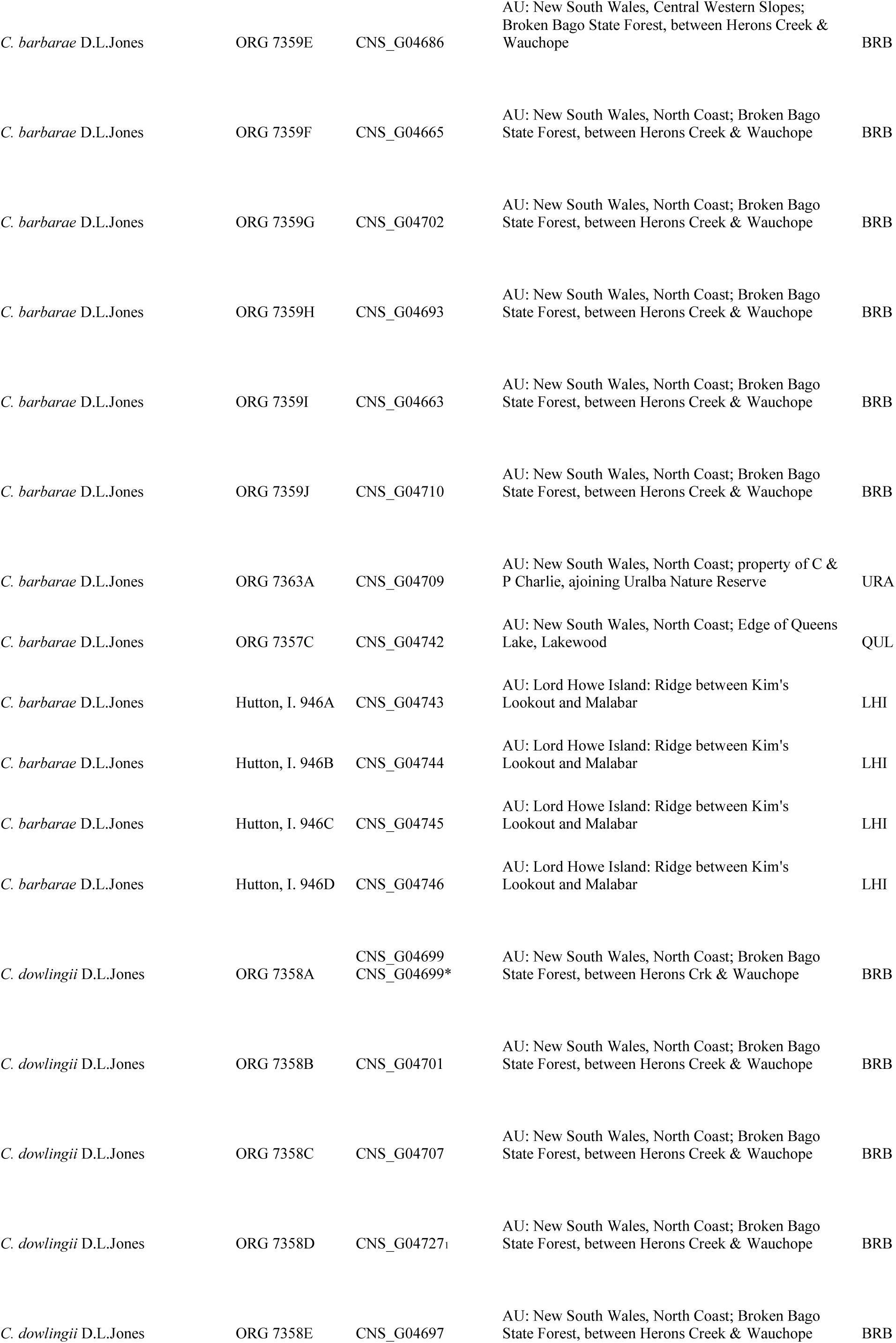
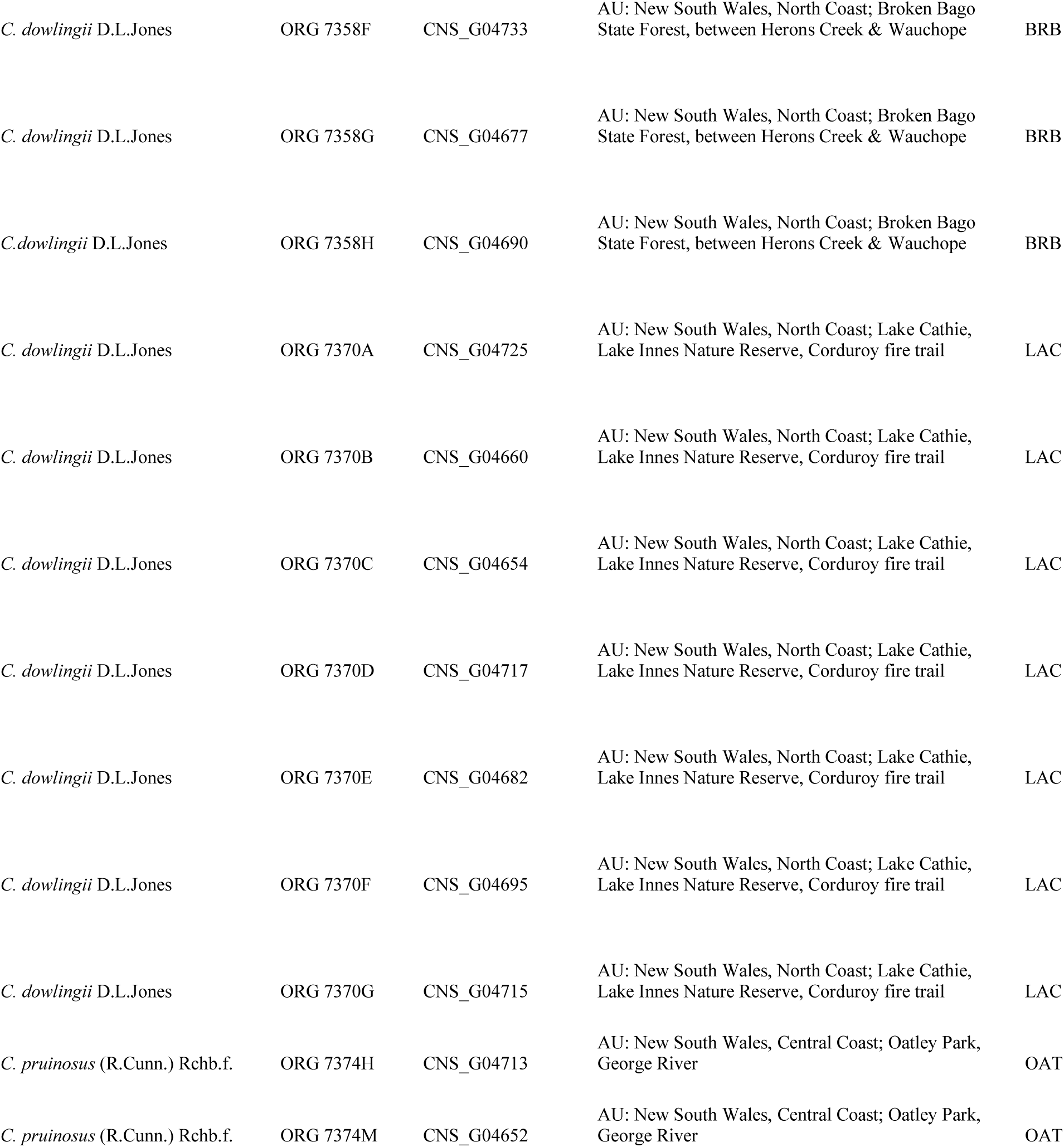
Material studied. AU: Australia; CNS: Australian Tropical Herbarium; ORG: Orchid Research Group, Centre for Australian Plant Biodiversity Research, Canberra; ^1^sample used for ddRADseq establishment phase; duplicate samples are marked with an asterisk

### DNA extraction, ddRAD library preparation, and sequencing

Total DNA was extracted from silica-dried leaf material using a modified CTAB protocol (Weising et al. 2005). DNA quality was assessed using a spectrophotometer (NanoDrop, Thermo Scientific) and gel electrophoresis on 2 % agarose gels, and DNA quantity was determined using a Qubit fluorometer (Thermo Fisher Scientific).

Double-digest restriction-site associated DNA (ddRAD) sequencing libraries were prepared following Peterson et al. (Peterson et al. 2012). During the initial establishment phase, double restriction enzyme digests of genomic DNA were carried out for three *Corybas* samples (*C. aconitiflorus* CNS_G04732*, C. barbarae* CNS_G04671, *C. dowlingii* CNS_G04727) testing eight different restriction enzyme combinations comprising a six base cutter (*PstI* or *EcoR1*) and a four base cutter (*MspI*, *HypCH4VI*, *MseI* or *NIaIII*). Digests were followed by ligation of barcoded adapters compatible with the restriction site overhang, bead purification, and amplification of the non-size selected sequencing library via PCR. Sequencing libraries were evaluated based on gel electrophoresis using TapeStation (Agilent Technologies, Santa Clara, CA, USA) to select the most suitable restriction enzyme combination that indicated the least amount of likely repetitive sequences. After this initial test, the enzyme combination *Pst*I and *Nla*III was selected for the three *Corybas* samples, including ligation of barcoded adapters, purification of the pooled digested-ligated fragments followed by size selection via Blue Pippin (Sage Science, Beverly, Massachusetts, USA) for two ranges, a narrow (280-342bp) and a wide (280-375bp) range. The two pooled libraries were amplified via PCR with indexed primers, and sequenced on the Illumina MiSeq platform for single-ended, 160 bp reads at the Australian Genome Research Facility (AGRF; Melbourne, Victoria, Australia). The sequencing data for the narrow and wide size selected libraries was analysed using the Stacks pipeline (Catchen et al. 2011, 2013) to assess the number of ddRAD loci per sample, the average coverage per sample, and the number of unique and shared ddRAD loci across the three samples. The narrow library yielded between 116,991 and 276,091 ddRAD loci per sample and an average coverage of 6.5-9.2 per sample and the wide library resulted in 85,229 to 203,174 ddRAD loci per sample and an average coverage of 6.4–9.4 per sample. The narrow size selection yielded a higher number of shared ddRAD loci across the three samples than the wider size selected sequencing library. Based on the evaluation of two pooled libraries, the narrow size selection was chosen for ddRAD sequencing for the complete sample set. Quality and reproducibility of libraries and DNA sequencing were assessed by running five samples in duplicate (6.9 % of all samples). Multiplexed libraries were sequenced on one lane of a NextSeq500 sequencing platform (Illumina Inc., San Diego, CA, USA) as single-ended, 150 bp reads at the Australian Genome Research Facility (AGRF; Melbourne, Victoria, Australia).

### Bioinformatics and data filtering

Quality of the sequence reads was examined using FastQC v.0.11.5 (Andrews 2010). Raw sequences were demultiplexed, trimmed and further processed using the ipyrad pipeline v.0.6.15 (Eaton and Overcast 2016). In an initial filtering step, reads with more than five low quality bases (Phred quality score < 20) were excluded from the data set. The phred quality score offset was set to 33. The strict adapter trimming option was selected and a minimum read length of 35bp after trimming was chosen to retain a read in the dataset. After these quality-filtering steps, the reads were clustered within and across samples by similarity of 85% using the vclust function in VSEARCH (Edgar 2010). The alignment was carried out using MUSCLE (Edgar 2004) as implemented in ipyrad. Clusters with less than six reads were excluded in order to ensure accurate base calls. The resulting clusters represent putative RAD loci shared across samples. A maximum number of five uncalled bases (‘Ns’) and a maximum number of eight heterozygote sites (‘Hs’) was allowed in the consensus sequences. The maximum number of SNPs within a locus was set to ten and the maximum number of indels per locus to five. For the sample set including all accessions of the *C. aconitiflorus* complex as well as two accessions of *C. pruinosus* as outgroup ipyrad runs for two different datasets were performed, i.e. based on loci shared by 20 individuals (m20) and on loci shared by 70 individuals (m70). Additionally, the same settings were used for ipyrad runs excluding the outgroup (*C. pruinosus,* 2 samples). The datasets generated and analysed during the current study are available at CSIRO’s Data Access Portal, [*DOI will be provided in published manuscript*].

### Phylogenomic and genetic structure analysis

Phylogenetic relationships were inferred using maximum likelihood (ML) based on concatenated alignments applying the GTR+ Γ model of nucleotide substitution using RAXML v.8.2.4 (Stamatakis 2014) for both datasets (m20, m70) including the outgroup. Statistical support was assessed via a rapid bootstrapping with 100 pseudoreplicates (Stamatakis et al. 2008)(Stamatakis et al. 2008) under the same ML analysis settings.

NeighborNet analysis of the *C. aconitiflorus* complex was carried out in SplitsTree v.4.13 (Huson and Bryant 2006) using the unlinked SNP output of the m20dataset excluding the outgroup. The unlinked SNPs represent one randomly chosen SNP per locus and are therefore considered independend markers. Equal angle split transformation and uncorrelated P distance were selected for the NeighborNet analysis.

Genetic structure of the *C. aconitiflorus* complex was analysed using the Bayesian Markov Chain Monte Carlo (MCMC) clustering method implemented in the program Structure v.2.3.4 (Pritchard et al. 2000) based on both datasets (m20, m70) excluding the outgroup. For both datasets, the Structure output format of unlinked SNPs (.ustr) of the ipyrad pipeline was used as input file. Data analysis assumed correlated allele frequencies and admixture and prior population information was not included in the analysis (Hubisz et al. 2009). After preliminary runs with a smaller number of cycles and because of computational limitations, we conducted three independent runs for each value of K = 3–8 with 100,000 MCMC cycles following a burn-in of 10,000 MCMC cycles for the final analyses. The range of K-values was chosen based on the number of expected species (3) and number of eight observed clades in the RAxML analysis. The number of genetic groups best fitting the dataset was determined using the delta K method (Evanno et al. 2005) as implemented in Structure Harvester (Earl and vonHoldt 2012).

## Results

An average of 2,98 (± 1,01) filtered Illumina reads per sample were used for the analyses. A total number of 362,412 pre-filtered loci passed the ipyrad pipeline. After subsequent filtering steps, the number of retained loci for the final datasets varied between 3,597 (m70) and 12,420 (m20) for the datasets including the outgroup, and between 4,293 (m70) and 14,915 (m20) loci for the datasets excluding the outgroup. The latter included 60,489 SNPs (see Table 2). The average read depth per locus was 20.87 (±4.01) reads. Further statistics of the ddRAD datasets are summarized in Tab. 2.

**Tab. 2.** Statistics for the ddRAD dataset for the *C. aconitiflorus* complex a) including the outgroup (*C. pruinosus*, 2 samples) and b) excluding the outgroup resulting from the different filtering thresholds in the ipyrad pipeline for loci shared by minimum number of samples. pis: parsimony informative characters, bp: base pairs, m: minimum number of samples

| Filtering threshold | m20 | m70 |
| --- | --- | --- |
| #RAD loci | 12,420 | 3,597 |
| #variable sites | 54,177 | 17,378 |
| #pis | 36,486 | 12,302 |
| #aligned bp | 1,682,315 | 487,474 |
| missing data* | 37.66% | 4.55% |

| Filtering threshold | m20 | m70 |
| --- | --- | --- |
| #RAD loci | 14,915 | 4,293 |
| #variable sites | 60,489 | 18,828 |
| #pis | 39,470 | 12,252 |
| #unlinked SNPs | 13,708 | 4,066 |
| #aligned bp | 1,967,151 | 581,128 |
| missing data* | 36.79% | 2.30% |

\*Proportion of gaps and undetermined bases in DNA sequence alignment

### Maximum likelihood analysis

The ML tree reconstruction based on the m20 dataset including the outgroup retrieved the *C. aconitiflorus* complex as monophyletic group with maximum bootstrap support (BS 100) (Fig. 2). Genetic divergence between the *C. aconitiflorus* complex and *C. pruinosus* was considerably higher than within species as indicated by branch length (Fig. 2). Within the C. aconitiflorus complex, the ML reconstruction did not provide support for the monophyly of the three species *C. aconitiflorus*, *C. barbarae,* and *C. dowlingii* (Fig. 2).

**Fig. 2.**
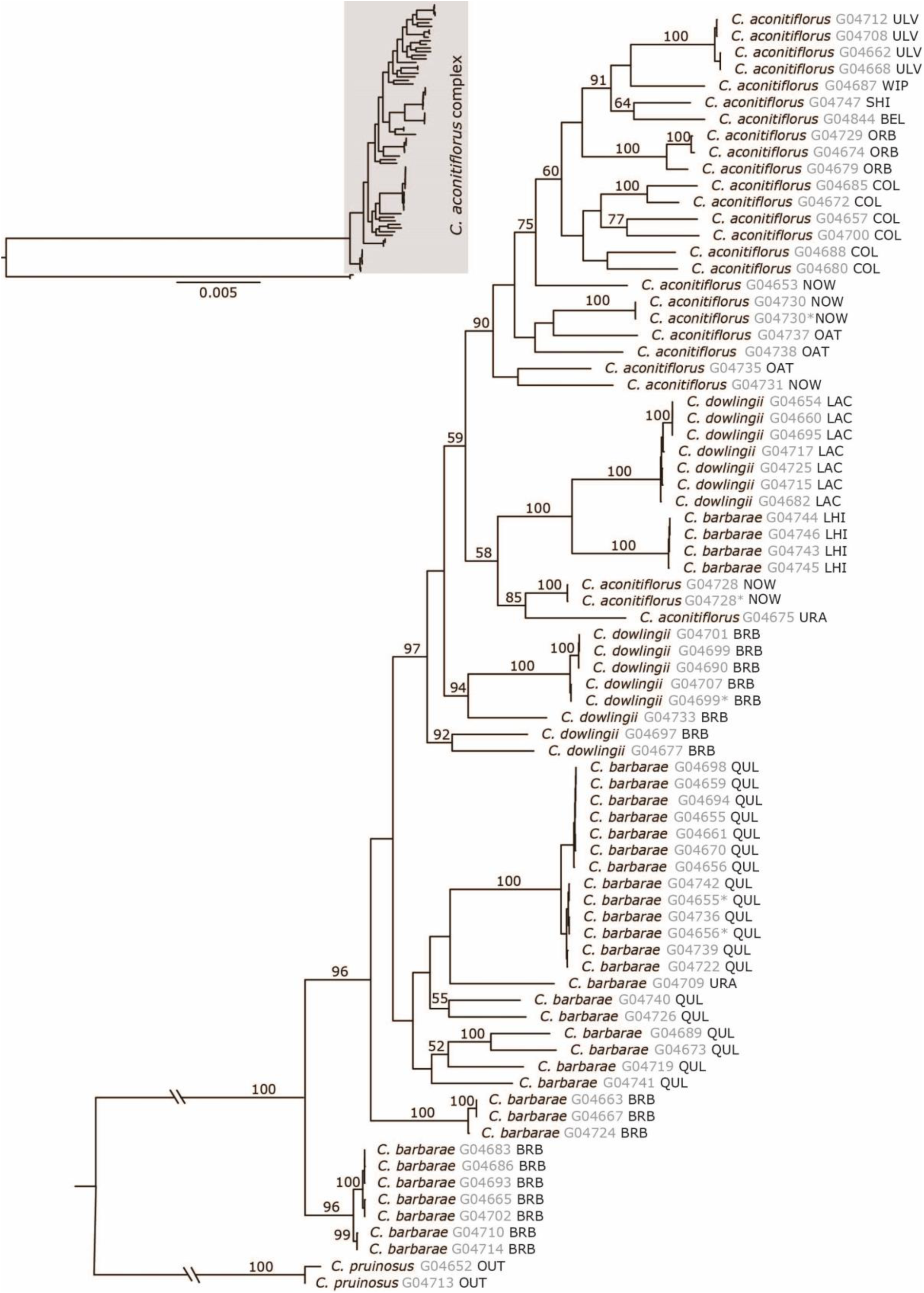
Maximum likelihood phylogeny of the *Corybas aconitiflorus* complex based on 12,420 ddRAD loci (m20 dataset including outgroup). BEL: Belgrave South; BRB: Broken Bago; COL: Colquhoun; LAC: Lake Cathie; LHI: Lord Howe Island; NOW: Nowra; OAT: Oatly Park; ORB: Orbost; QUL: Queens Lake; SHI: Shallow Inlet; URA: Uralba Nature Reserve; ULV: Ulverstone; WIP: Wilson Promontory. Bootstrap support values above 50 are given above branches. Duplicate samples are marked with an asterisk

A well-supported cluster (BS 96) comprising individuals from *C. barbarae* from North Coast (Broken Bago) was depicted as branching off first, followed by a second well supported cluster of *C. barbarae* (BS 100) from North Coast (Broken Bago). The next dichotomy showed one branch unifying individuals from *C. barbarae* from North Coast (Queens Lake and Uralba) and a highly supported branch (BS 97) harbouring individuals from *C. barbarae*, *C. dowlingii* and *C. aconitiflorus*. Within the latter branch, the first two diverging clusters were formed by individuals of *C. dowlingii* from North Coast (Broken Bago), both receiving high support (BS 92 and BS 94, respectively). The next diverging branch showed a dichotomy with one branch that harboured a moderately supported cluster (BS 85) unifying individuals of *C. aconitiflorus* from North Coast and South Coast, respectively (Uralba and Nowra) next to a highly supported cluster (BS 100) comprising individuals of *C. barbarae* and *C. dowlingii*. The latter cluster split into two highly supported clusters (BS 100), one with all individuals of *C. barbarae* from Lord Howe Island and the other with individuals of *C. dowlingii* from North Coast (Lake Cathie).

The second branch of the dichotomy was well supported (BS 90) and comprised the remaining samples of *C. aconitiflorus* from Central Coast and South Coast (Sydney and Nowra), all *C. aconitiflorus* samples from south Victoria (Colquhoun, Belgrave, Orbost, Shallow Inlet, Wilson Promontory) and Tasmania (Ulverstone) (Fig. 2). Within the latter, the *C. aconitiflorus* individuals from south Victoria and Tasmania formed a weakly supported subcluster (BS 60).

The ML analysis of the m70 dataset yielded congruent results for highly supported relationships and differed in topology for nodes that remained unsupported or received low statistical support in the analysis. Results of the ML analysis of the m70 dataset are presented in Online Resource 1.

### NeighborNet analysis

The NeighborNet network based on 13,708 unlinked SNPs of the m20 dataset of the *C. aconitiflorus* complex showed one large group comprising *C. barbarae* from the Australian mainland, one large group comprising *C. aconitiflorus,* and in intermediate position within the NeighborNet *C. barbarae* from Lorde Howe Island and *C. dowlingii* (Fig. 3). However, genetic distances between the branches in the NeighborNet were overall low and exhibited patterns of reticulation. Subgroups within the network were largely consistent with well supported clusters found in the ML analysis.

**Fig. 3.**
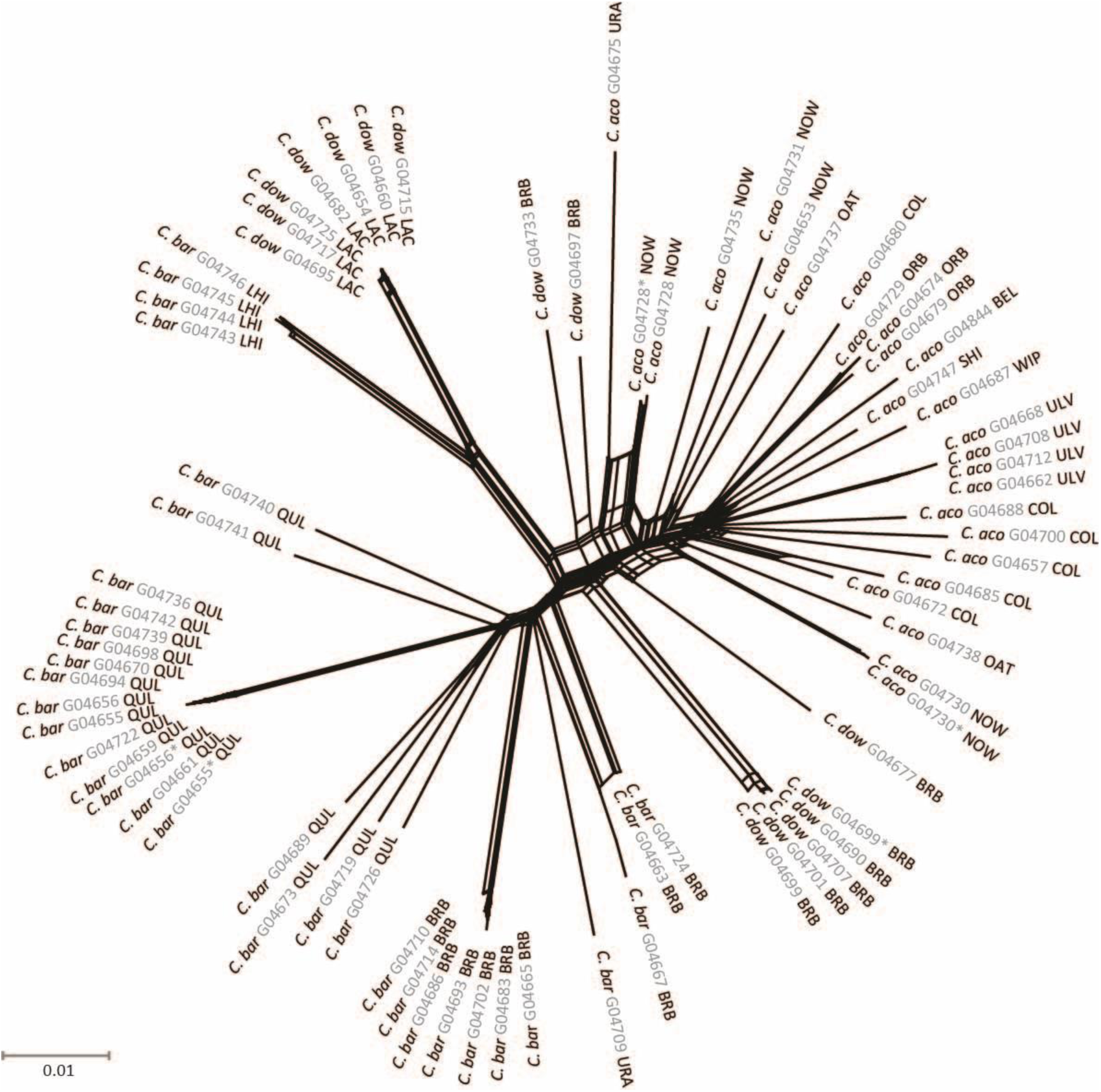
NeighborNet network for *Corybas aconitiflorus* complex based on 4,293 unlinked SNPs (m20 dataset excluding outgroup). C. aco: *Corybas aconitiflorus*; C. bar: *Corybas barbarae*; C. dow: *Corybas dowlingii*. BEL: Belgrave South; BRB: Broken Bago; COL: Colquhoun; LAC: Lake Cathie; LHI: Lord Howe Island; NOW: Nowra; OAT: Oatly Park; ORB: Orbost; QUL: Queens Lake; SHI: Shallow Inlet; URA: Uralba Nature Reserve; ULV: Ulverstone; WIP: Wilson Promontory. Duplicate samples are marked with an asterisk

Within the large group of *C. barbarae* from the Australian mainland, the NeighborNet diagram showed three branches that harboured individuals from Queens Lake (North Coast), two branches that comprised individuals from Broken Bago (North Coast) and one branch with *C. barbarae* from North Coast (Uralba). *Corybas barbarae* samples from Lord Howe Island formed a group that clustered together with *C. dowlingii* from Lake Cathie (North Coast), which corresponded to the relationships found in the ML analysis. Within the *C. aconitiflorus* group, samples from south Victoria and Tasmania formed a weakly differentiated subgroup, whereas *C. aconitiflorus* samples from NSW (Nowra, Sydney, Uralba) were found in more proximate position to *C. dowlingii* from Broken Bago, corresponding to relationships found in the ML analysis (Fig. 3). The NeigbourNet analysis based on the m70 dataset yielded highly congruent results to the analysis of the m20 dataset and are provided in Online Resource 2.

### Genetic structure analysis

In the following we report on the results of the Bayesian cluster analysis with Structure based on the m70 SNP dataset, which comprised 4,066 unlinked SNPs and a lower proportion of missing data than the m20 dataset as missing data can introduce biases to Structure analyses. For the m70 dataset, the best number of genetic groups (K) as determined by the modal Δ K distribution was K=7. The resulting bar plot (Fig. 4) showed several distinct genetic clusters and samples with varying degrees of genetic admixture. The genetic clusters retrieved by the Structure analysis did not correspond to current species concepts in the *C. aconitiflorus* complex.

**Fig. 4.**
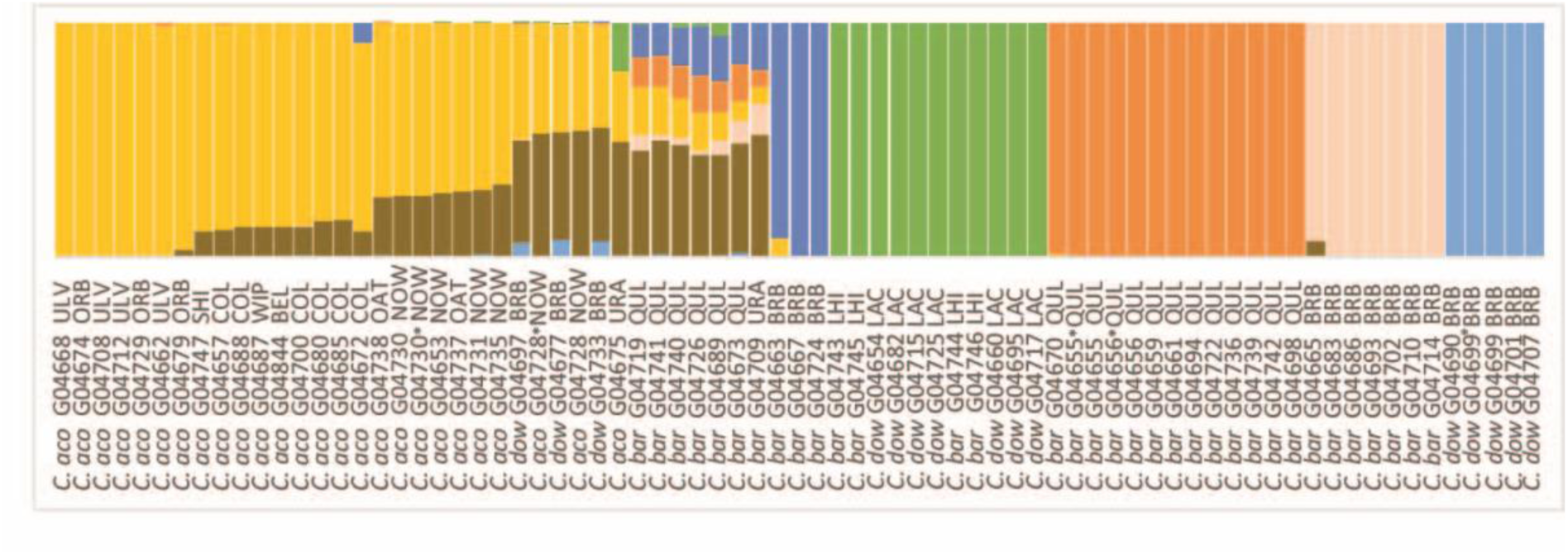
Genetic structure of *Corybas aconitiflorus* complex based on Structure analysis of 13,708 unlinked SNPs (m70 dataset excluding outgroup) for the optimal K value of seven genetic clusters. C. aco: *Corybas aconitiflorus*; C. bar: *Corybas barbarae*; C. dow: *Corybas dowlingii*. BEL: Belgrave South; BRB: Broken Bago; COL: Colquhoun; LAC: Lake Cathie; LHI: Lord Howe Island; NOW: Nowra; OAT: Oatly Park; ORB: Orbost; QUL: Queens Lake; SHI: Shallow Inlet; URA: Uralba Nature Reserve; ULV: Ulverstone; WIP: Wilson Promontory. Duplicate samples are marked with an asterisk

The majority of *C. barbarae* samples from the mainland fell into three genetically distinct clusters that exhibited a low or no signal of genetic admixture. Two of these clusters unified samples from North Coast (Broken Bago), which also formed two well-supported clades in the ML analysis and two distinct branches in the NeighborNet analysis. The third *C. barbarae* cluster in the Structure analysis comprised a group of eleven samples from North Coast (Queens Lake), which were also found as highly supported cluster in the ML analysis and as a distinct branch in the NeighborNet analysis. The *C. barbarae* samples from Lord Howe Island and the *C. dowlingii* samples from Lake Cathie together formed another genetically distinct cluster. These samples were also found as highly supported clade in the ML analysis and in the NeighborNet analysis. For the remaining *C. barbarae* samples from North Coast (Queens Lake) and from North Coast (Uralba) the Structure analysis indicated high levels of genetic admixture (Fig. 4). High levels of genetic admixture were also found for *C. dowlingii* from Broken Bago (North Coast) and several samples of *C. aconitiflorus* from South Coast (Nowra). The remaining *C. aconitiflorus* samples from Central Coast and South Coast (Sydney and Nowra) displayed moderate levels of genetic admixture, whereas the *C. aconitiflorus* samples from south Victoria and Tasmania displayed low levels or no genetic admixture in the Structure bar plot (Fig. 4). The results from the Structure analysis based on the m20 dataset are provided in Online Resource 3.

## Discussion

Accurate delimitation of species and infraspecific taxa is of critical importance to conservation biology as these are commonly used measures of biodiversity (Coates et al. 2018). However, taxon delimitation still heavily relies on the evaluation of morphological and ecological traits, which can be subject to convergent or parallel evolution and environmental plasticity. Further, taxonomic boundaries can be blurred through hybridisation and introgression, which are frequently observed phenomena among closely related orchid species (Dressler 1981; Cozzolino et al. 2006; Pinheiro et al. 2010; Nauheimer et al. 2018).

Recent advances in high throughput sequencing and statistical analyses now facilitate unprecedented fine-scale resolution of inter- and intraspecific relationships with detailed insights into genetic diversity and population structure. Hence, these approaches provide critical genomic data to evaluate taxonomic concepts in species complexes and to inform the development of effective conservation strategies for threatened species. Reduced representation library sequencing approaches, such as RADseq, ddRADseq and genotyping-by-sequencing (GBS), have proven to be powerful tools for disentangling complex relationships and delimiting taxonomic boundaries in orchids (Ahrens et al. 2017; Bateman et al. 2018; Brandrud et al. 2019a). Here, we present a study using more than 13,000 SNPs to unravel the complex relationships in the *C. aconitiflorus* complex containing three morphologically similar orchid species endemic to Australia.

Our genomic study provided insights into genetic diversity and structure of the *C. aconitiflorus* complex at fine-scale resolution. The results from the molecular analyses present a complex pattern that indicates occasional gene flow between *C. aconitiflorus* and *C. barbarae*. The apparent non-monophyly of the three species in the phylogenetic results is coherent with patterns expected in the presence of reticulation. The phylogenetic reconstruction showed genetically admixed samples of *C. aconitiflorus* and *C. barbarae* either towards the basal position of the main clade for each species comprising samples which exhibited no or low levels of admixture, or in intermediate position between the *C. aconitiflorus* and *C. barbarae* main clades. In the phylogenetic tree reconstruction, *Corybas dowlingii* clustered most closely to samples of *C. aconitiflorus* and *C. barbarae* for which genetic signatures of hybridisation were detected in the NeigborNet and Structure analysis.

The NeighborNet and Structure analyses revealed conflicting phylogenetic signal and genetic admixture in several samples, providing evidence for occasional hybridisation and introgression between *C. aconitiflorus* and *C. barbarae*. In the NeighborNet analysis (Fig. 3), the majority of *C. aconitiflorus* and *C. barbarae* samples formed a cluster each, while all *C. dowlingii* samples were found in intermediate position, together with several *C. aconitiflorus* and *C. barbarae* samples. Further, the NeighborNet analyses indicated conflicting genetic signal for the latter *C. aconitiflorus*, *C. barbarae* and *C. dowlingii* samples, which is consistent with patterns expected in the presence of hybridisation and introgression. The Structure analysis revealed considerable genetic structure within the species complex with seven genetic groups, and detected genetic admixture in several *C. aconitiflorus*, *C. barbarae* and *C. dowlingii* samples (Fig. 4). Within *C. aconitiflorus*, the highest degrees of genetic admixture were found in areas where the distributional range overlaps with *C. barbarae*, whereas *C. aconitiflorus* populations from areas where *C. barbarae* was absent (i.e. in Ulverstone in Tasmania and five locations in Victoria) showed no or low levels of genetic admixture. Thus, the results of this study indicate occasional hybridisation and introgression between *C. barbarae* and *C. aconitiflorus,* and provide genetic evidence that *C. dowlingii* is of hybrid origin. However, the genetic results also showed that samples of *C. aconitiflorus* and *C. barbarae*, respectively, are genetically more similar than to the other species, with the exception of samples which exhibited genetic signatures of hybridisation. Notably, *C. aconitiflorus* and *C. barbarae* samples which did not show signatures of admixtures genetically distinct from the other species in all analyses. Furthermore, genetically pure samples from different populations of the same species consistently clustered more closely together than those from co-occurring populations of the other species. This indicates that the two species, *C. aconitflorus* and *C. barbarae* maintain genetic coherence despite occasional hybridisation in areas where they co-occur.

The detected signatures of hybridisation and introgression are also in line with the presence of intermediate morphological character states in this species complex, in particular in floral traits such as flower colour and size. The intermediate position of *C. dowlingii* in the NeighborNet graph as well as the clustering of *C. dowlingii* samples with *C. barbarae* samples in the Structure analysis and the signature of genetic admixture found in several *C. aconitiflorus, C. dowlingii* and *C. barbarae* samples in the Structure analysis, point to a hybrid origin of *C. dowlingii*. Notably, the study also uncovered signatures of hybridisation and introgression in individuals that did not exhibit the *C. dowlingii* morphotype and which were morphologically assigned to either *C. aconitiflorus* or *C. barbarae.* These results highlight existing challenges in interpreting morphological variation in the complex and thus in identifying hybrids and introgressed individuals based on morphological traits alone.

Based on the results of this genomic study, we propose to change the taxonomic status of *C. dowlingii* in accordance with Article 50 of the International Code of Nomenclature for algae, fungi, and plants (Turland et al. 2018) to reflect its hybrid origin, as follows: *Corybas × dowlingii* D.L.Jones, The Orchadian 14(9): 419-420, f.1 (2004) (pro sp.).

### Assessment of conservation status of *C. × dowlingii*

Based on the molecular results of this study, which indicate that *C. × dowlingii* is of hybrid origin, the question arises whether and to what extent the taxon may merit conservation. A recent review of conservation legislation found that legal definitions of species are quite flexible and can accommodate a range of infra-specific taxa and divergent populations (Coates et al. 2018). While traditionally the protection of pure, genetically distinct species that do not interbreed successfully have been favoured in conservation science and policy (Agapow et al. 2004), it is increasingly recognised that biological diversity generated by hybridisation can also hold conservation value (Allendorf et al. 2001; Agapow et al. 2004).

While the evolutionary importance of hybridisation in the diversification of plants has long been recognised, recent genomic studies highlighted the prevalence of hybridisation in the natural world (Taylor and Larson 2019). Further, the detection of ancient hybridisation in genomic studies, which indicate that hybridisation occurred in many taxa at some point in the past, has led to a greater appreciation of the evolutionary importance of hybridisation and introgression and improves our understanding of potential long-term consequences of hybridisation (Taylor and Larson 2019). Hybridisation can lead to greater fitness compared to parental species, i.e. heterosis (hybrid vigour), and is seen as important evolutionary processes that promotes adaptation and speciation. Hybrids can exhibit novel adaptive traits that allow for increased ecosystem resilience to environmental stressors and can allow for the successful colonisation of novel habitats (Stebbins 1959). However, hybrids can also have negative impacts on biodiversity, in particular in cases where hybrids pose risks to the survival of their parental species or to other native vegetation (Rhymer and Simberloff 1996). Based on a review of species hybrids and their conservation or management, Jackiw et al. (Jackiw et al. 2015) strongly advocate for a case by case approach in assessing the conservation value of hybrids.

In the following, we will apply the framework to guide the conservation of species hybrids based on ethical and ecological considerations developed by Jackiw et al. (Jackiw et al. 2015) to assess the conservation value of *C. × dowlingii*. A key ecological consideration is whether the hybrid is likely to pose a risk to the survival of the parental species, for example through decreasing the genetic variability in parental species or by causing extinction through genetic assimilation as long-term consequences of recurrent hybridisation and introgression (Rhymer and Simberloff 1996). This is of particular importance in cases where parental species are already threatened by other factors. *Corybas × dowlingii* has only a narrow distribution whereas both parental species are common and occur over a large distributional range with both sympatric and allopatric distributions. From this perspective, the hybrid species *C. × dowlingii* as well as hybrids that occasionally form between the two parental species and which do not exhibit the *C. dowlingii* morphotype, are unlikely to negatively affect the long-term evolutionary persistence of the parental species.

Another consideration concerns species fitness as hybrid species exhibiting a higher fitness than the parental species may have detrimental effects either for the parental species or to other native species, as has been documented for example in invasive weeds (Stebbins 1959; Rhymer and Simberloff 1996). *Corybas × dowlingii* is only known from a few populations extending ca. 100 km from Bulahdelah to Freemans Waterhole in New South Wales, overlapping with the geographic distribution of the two parental species and occurring in the same habitat. Based on our ecological observations in the field, there is no indication of increased fitness of *C. × dowlingii* compared to its parental species. From this perspective, the hybrid species is also unlikely to negatively affect the survival of the parental species or other native vegetation.

An important consideration from an evolutionary perspective is that hybrids are often regarded as beneficial. They hold the potential to act as catalysts for speciation or more generally as pathway for evolution, for example in cases where hybrids exhibit novel properties such as in floral scent (Stökl et al. 2008; Vereecken et al. 2010), habitat requirements (Jacquemyn et al. 2012) or genome duplication and rearrangements (de Storme and Mason 2014). These novel traits might lead subsequently to the formation of a new species. Therefore, the maintenance of evolutionary processes becomes a key ethical consideration in the conservation of species hybrids (Jackiw et al. 2015). Given that *C. × dowlingii* is unlikely to pose a threat to the long-term survival of its parental species nor other native species, in the following we will examine possible beneficial aspects that may warrant the conservation of *C. × dowlingii,* such as the potential of the hybrid species to act as pathway for evolution.

In the New Caledonian hybrid species *Corybas × halleanus* E.Faria, a greater tolerance to lower humidity was observed for the hybrid species compared to its more moisture dependent parental species (Faria 2016). The hybrid species thus exhibits increased resilience to environmental stressors, which enables the hybrid to colonise novel habitat, thus rendering the conservation of this hybrid species beneficial. In contrast, *C. × dowlingii* is found in the same habitat as its parental species and growing sympatrically with *C. barbarae* (Jones 2004), thus not exhibiting novel habitat requirements.

In terms of pollination biology, observations on pollinators in the *C. aconitiflorus* complex are still scarce. While the pollination strategy of the genus has been regarded as food or brood deceptive based on observations from other *Corybas* species (Pridgeon et al. 2001), recent observations in *C. aconitiflorus* indicate that the species is food-rewarding (Kuiter and Findlater-Smith 2017). Fungus gnats of the genus *Phthinia* Winnertz 1863 (Mycetophilidae) were observed to visit the flowers to forage on the column mound which has been reported to exude nectar and females where found leaving the flowers with pollinia attached to their thorax (Kuiter and Findlater-Smith 2017). The repeated and directed visiting behaviour of the fungus gnats was regarded as indication that the fungus gnats are attracted through floral scent. Further, the plant-pollinator relationship was found to be specific in the *Corybas* populations in Victoria, where only one *Phthinia* species was observed to visit the flowers (Kuiter and Findlater-Smith 2017). However, the detection of genetically admixed individuals in our molecular study implies that the plant-pollinator relationship between *C. aconitiflorus* and *C. barbarae* is less specific and allows for occasional cross-pollination.

Orchid hybrids hold the potential to develop novel floral traits, which can result in reproductive isolation between the hybrid and parental species. This is of particular importance in orchids as the plant-pollinator interaction is often highly specific (Nilsson 1992). For example, the hybrid between two European *Ophrys* species was found to exhibit a novel floral scent which resulted in a pollinator shift (Vereecken et al. 2010). In the sun orchids (*Thelymitra*), hybrids are often found which have shifted their pollination system from outcrossing to obligatorily selfing (Jeanes 2004, 2009, 2013), which likewise presents an immediate reproductive isolation mechanism. Hence, pollination studies in *C. × dowlingii* are desirable to assess if the hybrid exhibits novel floral traits leading to reproductive isolation from its parental species. Likewise, changes in ploidy levels in association with hybridisation (allopolyploidisation) can also lead to immediate reproductive isolation between the hybrid and its parental species. Allopolyploid hybrid species tend to derive from parents that are evolutionarily more divergent than parents of homoploid hybrid species (Chapman and Burke 2007; Paun et al. 2009). In *Corybas*, cytogenetic studies found evidence for polyploidisation in the genus, e.g. in *C. cheesmanii* (Hook.f.) Kuntze with 2=54+2, pointing to a triploid or hexaploid origin (Dawson et al. 2007), however no chromosome counts are available for the *C. × dowlingii* nor its parental species. To assess possible reproductive isolation of *C. × dowlingii* from its parental species based on genome duplication or re-arrangements, cytogenetic studies in the *C. aconitiflorus* complex are required. However, given that hybridsation between closely related species is more likely to give rise to homoploid hybrids (Chapman and Burke 2007; Paun et al. 2009), we consider the establishment of reproductive isolation of *C. × dowlingii* from its parental species through genome duplication or re-arrangements as unlikely.

Our overall assessment of key ecological and societal considerations indicated a low conservation concern for *C. × dowlingii* and highlighted areas for further studies to assess the degree of reproductive isolation of *C. × dowlingii* from its parental species to increase our understanding of the potential evolutionary fate of this species. In an environment of limited financial resources, it is of key importance to consider whether it is ethical to direct conservation funds and effort to hybrid species. In the case of *C. × dowlingii*, our overall assessment under the current available evidence points toward a low priority for conservation of the hybrid species, in particular considering that the two parental species are common and tend to hybridise occasionally. Hence, the maintenance of evolutionary processes that may result in the emergence of novel traits driving further diversification appears warranted in this species complex without particular conservation emphasis on the hybrid species *C. × dowlingii*. Based on the results of this study, we regard the conservation status of *C. × dowlingii* of ‘least concern’ and recommend changing its current conservation status in the threatened species legislations of New South Wales and Australia accordingly.

## Conclusions

This conservation genomic study clarified taxonomic delimitation in the *C. aconitiflorus* complex and provided molecular evidence for occasional hybridisation and introgression between *C. aconitiflorus* and *C. barbarae*. *Corybas dowlingii* was found to be of hybrid origin and its taxonomic status was changed to *C. × dowlingii* to reflect this. Based on our assessment of the conservation status of *C. × dowlingii* considering key ecological and ethical aspects, we conclude that the conservation status of *C. × dowlingii* is of least concern and recommend this to be reflected in the Australian legislation at state and federal level.

## Supporting information

Online Resource 1

Online Resource 2

Online Resource 3

## Acknowledgments

The authors thank the Office of Environment and Heritage (Dept. of Planning and Environment, New South Wales, Australia) for financial support of the study and for granting of collection permits (NSW National Parks & Wildlife Service Scientific Licence SL100750 and Forest NSW Special Purpose Permit for Research XX51121). We thank R. Autin, C. Broadfield, J. Cootes, I. Hutton, W. Probert, J. Moye, J. Riley, A. Stephenson, R. Thomson and J. Turner for contributing plant material to the study. The authors acknowledge the use of lab services and sequencing facilities of the Australian Genomic Research Facility (AGRF).

## Online Resources

**Online Resource 1**. Maximum likelihood phylogeny of the *Corybas aconitiflorus* complex based on 3,597 ddRAD loci (m70 dataset including outgroup).

**Online Resource 2**. NeighborNet network for *Corybas aconitiflorus* complex based on 4,066 unlinked SNPs (m70 dataset excluding outgroup).

**Online Resource 3**. Genetic structure of *Corybas aconitiflorus* complex based on Structure analysis of 13,708 unlinked SNPs (m20 dataset excluding outgroup) for the optimal K value of seven genetic clusters.

