## Supplementary material for "Conservation genomics of an Australian orchid complex with implications for the taxonomic and conservation status of *Corybas dowlingii*": Online Resource 1

**Online Resource 1:** Maximum likelihood phylogeny of the *Corybas aconitiflorus* complex based on 3,597 ddRAD loci (m70 dataset including outgroup). BEL: Belgrave South; BRB: Broken Bago; COL: Colquhoun; LAC: Lake Cathie; LHI: Lord Howe Island; NOW: Nowra; OAT: Oatly Park; ORB: Orbost; QUL: Queens Lake; SHI: Shallow Inlet; URA: Uralba Nature Reserve; ULV: Ulverstone; WIP: Wilson Promontory. Bootstrap support values above 50 are given above branches. Duplicate samples are marked with an asterisk.

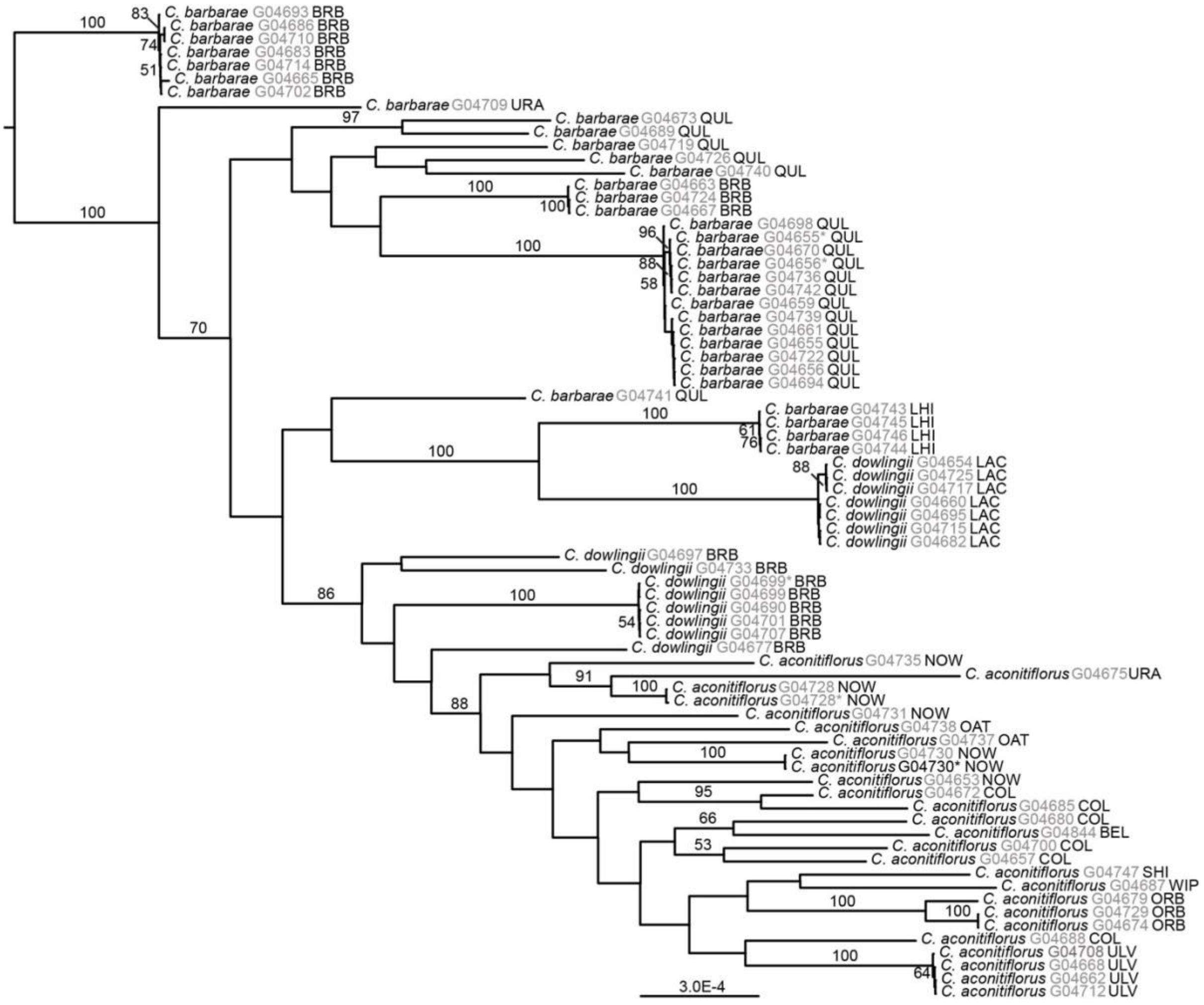
