## Supplementary material for "Conservation genomics of an Australian orchid complex with implications for the taxonomic and conservation status of *Corybas dowlingii*": Online Resource 3

**Online Resource 3:** Genetic structure of *Corybas aconitiflorus* complex based on Structure analysis of 13,708 unlinked SNPs (m20 dataset excluding outgroup) for the optimal K value of seven genetic clusters. C. aco: *Corybas aconitiflorus*; C. bar: *Corybas barbarae*; C. dow: *Corybas dowlingii*. BEL: Belgrave South; BRB: Broken Bago; COL: Colquhoun; LAC: Lake Cathie; LHI: Lord Howe Island; NOW: Nowra; OAT: Oatly Park; ORB: Orbost; QUL: Queens Lake; SHI: Shallow Inlet; URA: Uralba Nature Reserve; ULV: Ulverstone; WIP: Wilson Promontory. Duplicate samples are marked with an asterisk.

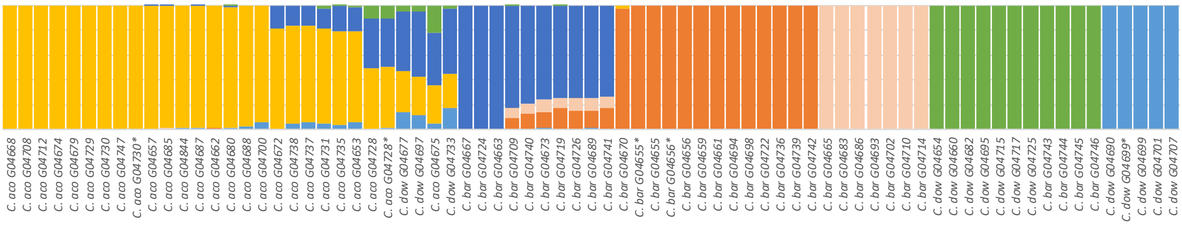
